# mSWI/SNF interacts with the ribosome and its inhibition/mutations alter translation and sensitize to mTOR/PI3K inhibitors

**DOI:** 10.1101/2021.05.10.443459

**Authors:** Livia Ulicna, Samuel C. Kimmey, Christopher M. Weber, Grace M. Allard, Sean C. Bedall, Gerald R. Crabtree, Gregory R. Bean, Capucine Van Rechem

## Abstract

The chromatin remodelers mammalian SWItch/Sucrose Non-Fermentable (mSWI/SNF) subunits are mutated, deleted or amplified in more than 40% of cancers. Understanding their functions in normal cells and the consequences of cancer’s alterations will lead to path toward new targeted therapies. Canonically, mSWI/SNF complexes regulate the structure of chromatin, however they likely have additional functions which could be relevant in carcinogenesis. Here, we highlight the substantial alteration of mSWI/SNF subunits expression in both the nucleus and cytoplasm in breast cancer cases. We demonstrate mSWI/SNF cytoplasmic localization and interaction with the translation initiation machinery. Short-term inhibition and depletion of specific subunits alter protein synthesis, implicating a direct role for these factors in translation. Inhibition and depletion of specific subunits increase sensitivity to mTOR-PI3K inhibitors, suggesting a potential therapeutic opportunity for diseases harboring mutations in these complexes. Indeed, *SMARCA4* pathogenic mutations decrease protein synthesis. Furthermore, taking advantage of the DepMap studies, we demonstrate cancer cells harboring mutations of specific mSWI/SNF subunits exhibit a genetic dependency on translation factors and are particularly sensitive to translation pathway inhibitors. In conclusion, we report an unexpected cytoplasmic role for mSWI/SNF in protein synthesis, suggesting potential new therapeutic opportunities for patients afflicted by cancers demonstrating alterations in its subunits.

**Statement of significance:** This study establishes direct functions for mSWI/SNF in protein synthesis. mSWI/SNF inhibition, depletion and cancer mutations alter translation and increase sensitivity to translation pathway inhibitors, illustrating the potential for new therapeutic strategies.

## Introduction

The mammalian SWI/SNF (mSWI/SNF, SWItch/Sucrose Non-Fermentable) complexes are evolutionary conserved ATP-dependent chromatin remodelers that modify chromatin architecture and thus influence gene expression (*1–3*). In human, there are three main complexes: BRG1/BRM Associated Factors (BAF), Polybromo-associated BAF (PBAF), and non-canonical BAF (ncBAF) (*4*). These complexes are composed of both common and specific subunits which are altered in more than 40% of cancers (*5*). Indeed, genes coding for mSWI/SNF are not only mutated in 20% (*1–3*), but also exhibit copy number variation in 30% of cancers (*5*). Many mutations result in loss-of-function and have been shown to not only drive disease progression but also facilitate resistance to therapy (*1–3, 6*). Alteration rates of specific subunits are disease-specific, suggesting context-dependent functions (*1–3, 5*).

While mSWI/SNF functions are mostly studied in the nucleus in the context of nucleosome positioning through direct interaction between mSWI/SNF and the nucleosome, few studies investigate the possibility of non-chromatin substrates for these complexes. In yeast, SWI/SNF evicts Silent information regulator Sir3p from the nucleosome, requiring a direct interaction between these proteins (*7*), and the catalytic subunit Brg1 (SMARCA4) activates Mec1 kinase by direct protein-protein interaction (*8*). In humans, BAF directly evicts Polycomb Repressive Complex 1 (PRC1) from the chromatin (*9*). Furthermore, mSWI/SNF subunits are not only present in the cell nucleus but can also shuttle between nucleus and cytoplasm (*10, 11*). Differential localization could be relevant in disease. For example, SMARCA4 is cytoplasmic in a subset of corticotroph adenomas (*12*), SMARCA2 nuclear localization correlates with better prognosis in lung cancer (*13*), and cytoplasmic ARID1B promotes oncogenesis and correlates with advanced pancreatic cancer (*14*). Together, these studies provide evidence for possible disease-associated functions of mSWI/SNF complexes outside of their direct roles on the nucleosome.

In this study, analysis of breast cancer patient tissues reveals a substantial alteration of nuclear and cytoplasmic expression of the ATPases SMARCA2-4 and the PBAF-specific subunits PB1, BRD7 and ARID2. We further demonstrate that cytoplasmic mSWI/SNF subunits interact with the translation machinery and affect protein synthesis. mSWI/SNF inhibition and depletion in combination with mTOR-PI3K inhibitors decrease cell viability, suggesting a possibility for protein synthesis targeted therapy in mSWI/SNF-perturbed diseases. Indeed, expression of *SMARCA4* loss-of-function pathogenic mutations detected in cancers decrease translation. Finally, analysis of the DepMap database unveiled mSWI/SNF-mutated cancer cell lines’ genetic dependency on translation factors and sensitivity to translation pathway inhibitors. Together, our data support a direct role for mSWI/SNF subunits in protein synthesis with the potential to develop into new therapeutic strategies.

## Results

### Breast cancer specimens predominantly gain SMARCA4 and lose SMARCA2 expression

Several mSWI/SNF subunits are observed in the cytoplasm in tumors and altered cellular localization could be useful markers (*12–14*). While mSWI/SNF subunits are primarily considered tumor suppressors in association with cancer, evidence has begun to shed light on their potential oncogenic roles. For example, SMARCA4 is unlikely to be a tumor suppressor in breast cancer (*15*). Because mis-expression or localization is not necessarily associated with genomic alteration, we evaluated mSWI/SNF subunits in breast cancer cases by immunohistochemistry. We utilized a tissue microarray composed of 46 cases including primary tumors and nodal and distant metastases.

We first focused on the mutually exclusive ATPases. SMARCA4 was very minimally expressed in normal breast epithelium, with occasional cells being nuclear positive (Fig. 1A and Supplementary Fig. S1A). Hence, we only observed gain of SMARCA4 expression in breast cancer. Remarkably, 51% of cases presented a gain in nuclear and cytoplasmic SMARCA4 and 18% a weak cytoplasmic staining that could be considered as background (a potential limitation of IHC). Notably, five cases (11%) harbored a *SMARCA4* mutation, three of which were associated with gain-of-expression by IHC (Supplementary Fig. S1B).

**Figure 1.**
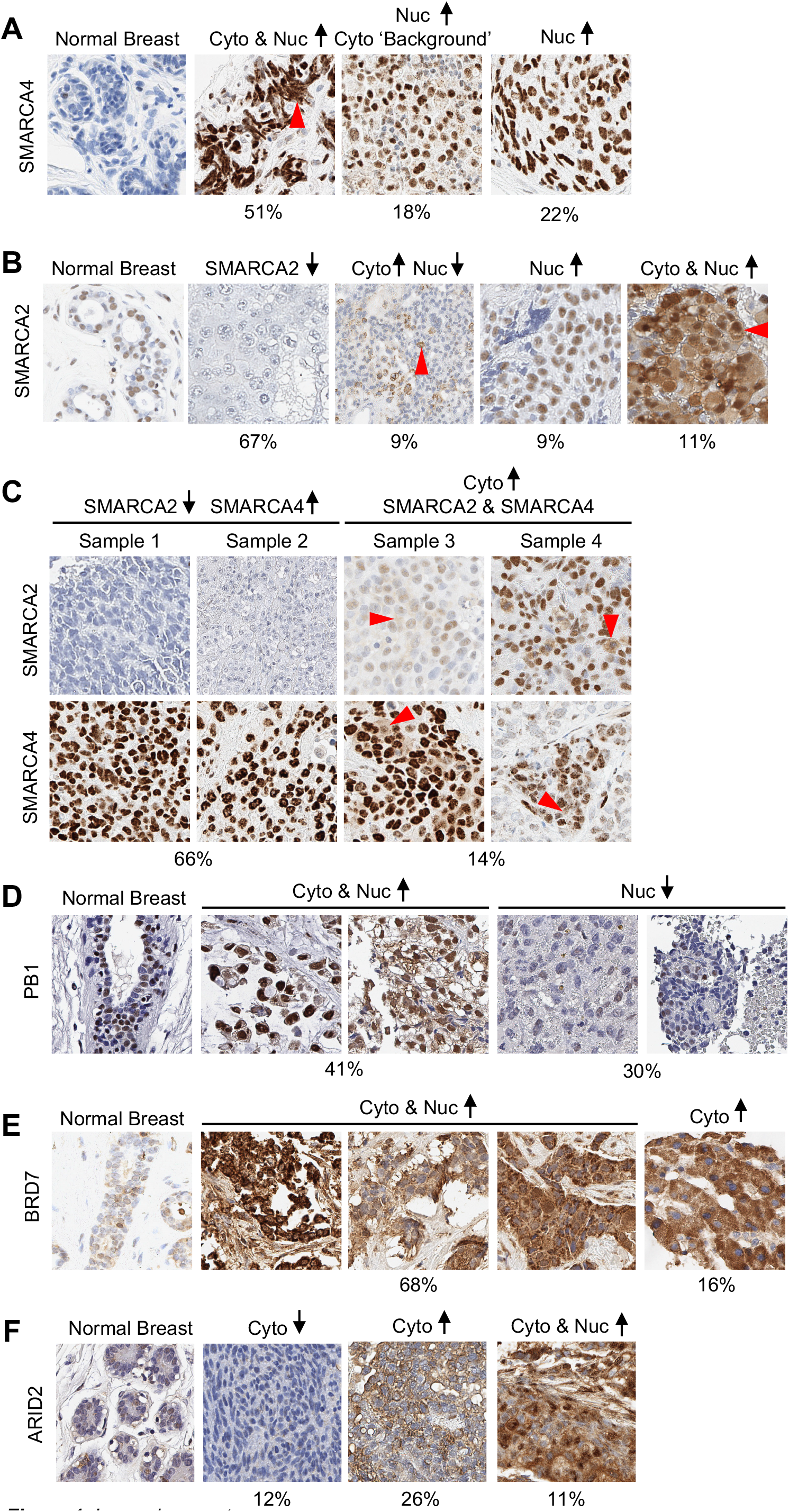
mSWI/SNF subunits expression and subcellular localization are altered in breast cancers. Representative examples of immunohistochemistry from breast cancer cases for SMARCA4 (**A**), SMARCA2 (**B**), identical tumors from the same patients for SMARCA2 and SMARCA4 (**C**), PB1 (**D**), BRD7 (**E**) and ARID2 (**F**). 20X images. Cyto = cytoplasmic, Nuc = nuclear.

SMARCA2 exhibited moderate nuclear expression in ~50% of normal breast epithelial cells (Fig. 1B and Supplementary Fig. S1C). 67% of cases demonstrated loss of nuclear SMARCA2 and 9% concomitantly gained cytoplasmic expression. Furthermore, 11% of cases showed increased nuclear and cytoplasmic SMARCA2 staining, and 7% exhibited gain of exclusive cytoplasmic staining. Of note, some cases presented both gains and losses of SMARCA2 expression within different samples.

Taken together, SMARCA2 is the primary mSWI/SNF ATPase expressed in normal breast and is predominantly lost in breast cancers. On the contrary, SMARCA4 is relatively overexpressed. Increased cytoplasmic staining of both ATPases was observed in 17% of cases. Gain of SMARCA4 with concurrent loss of SMARCA2 was predominant, with 66% of cases demonstrating a pattern of mSWI/SNF complexes supportive of a potential swap involving one ATPase for the other.

### PBAF-specific subunits expression is highly altered in breast cancers

Mutations in the PBAF-specific subunits *ARID2* and *PB1* are reportedly less frequent in breast cancers than the BAF-specific *ARID1A* and *ARID1B* (*2, 3*). Therefore, we decided to further investigate expression levels of PBAF subunits.

PB1 presented various levels of exclusive nuclear staining in normal breast epithelium (Fig. 1D and Supplementary Fig. S2A). Within our breast cancer cases, while 30% demonstrated a loss in nuclear staining, 41% showed substantial cytoplasmic expression and 22% exhibited weak cytoplasmic staining.

In normal breast, all cells showed cytoplasmic BRD7 with weak nuclear staining in less than 50% of the cells (Fig. 1E and Supplementary Fig. S2B). In breast cancer samples, BRD7 was prominently overexpressed: 68% of cases exhibited increased nuclear and cytoplasmic staining and 16% exhibited exclusively cytoplasmic gain.

Finally, ARID2 resembled BRD7 in normal breast, being predominantly cytoplasmic with few cells exhibiting a weak nuclear staining (Fig. 1F and Supplementary Fig. S2C). In breast cancer cases, 31% demonstrated gain in nuclear ARID2 expression, 26% a gain of exclusively cytoplasmic staining, 11% a gain in both compartments and 12% a loss of cytoplasmic ARID2.

All considered, while a substantial number of cases presented loss of expression, PBAF-specific subunits were primarily overexpressed in breast cancer cases. Of note, some patients’ tumors showed both gains and losses and/or variation in compartmental distribution across different samples, highlighting the complexity of tumor and cancer evolution observed in primary, recurrent and metastatic samples.

### Cytoplasmic mSWI/SNF subunits interact with the translation machinery

Because of the significant population of mSWI/SNF subunits in the cytoplasm in normal and breast cancer tissues, we further assessed their subcellular localization in cells harboring intact mSWI/SNF complexes (*4*). Biochemical fractionation from HEK293T revealed the substantial presence of BAF, PBAF and ncBAF subunits in the cytoplasmic compartment (Fig. 2A). Indeed, specific subunits are distributed equally in cytoplasmic and nuclear compartments (SMARCC1 and SMARCC2) and several are enriched in the cytoplasm compared to nucleus and chromatin combined (ARID2, PHF10, BRD7, ARID1A, ARID1B, DPF2 and BRD9). Based on the previously published role for the lysine demethylase KDM4A in the cytoplasm (*16*), we reasoned that cytoplasmic mSWI/SNF might be involved in protein synthesis. We performed polysome profiling and detected proteins present in collected fractions by western blot. All tested mSWI/SNF subunits were enriched in the initiating fractions (Fig. 2B-C and Supplementary Fig. S3). We further confirmed mSWI/SNF interaction with translation factors by coimmunoprecipitations (Fig. 2D). Treatment of extracts with RNAse A indicate that RNAs are likely not necessary for such interactions (Fig. 2D). Together, these data demonstrate mSWI/SNF’s interaction with the translation machinery and suggest that these complexes could directly impact translation.

**Figure 2.**
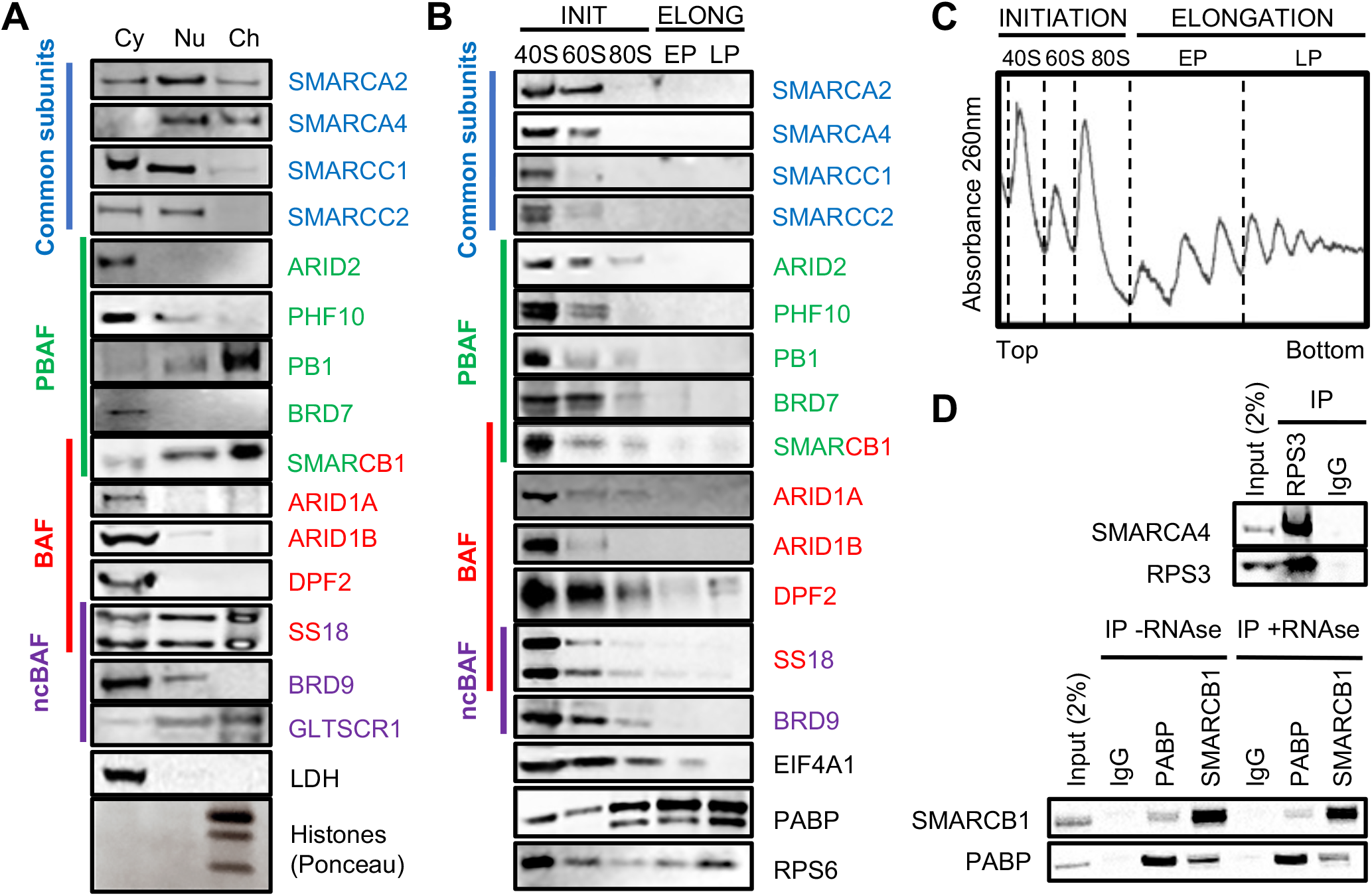
mSWI/SNF subunits localize in the cytoplasm and interact with the translation machinery. **A.** Immunoblot from fractionated HEK293T cells (Cy: cytoplasm, Nu: nucleus, Ch: chromatin). **B.** Immunoblots from HEK293T polysome profiles fractions (INIT: initiation, ELONG: elongation, EP: early polysomes, LP: late polysomes, as represented in C). **C.** Representative polysome profile from HEK293T cells. **D.** Coimmunoprecipitation from HEK293T cells. +RNAse: RNAse A was added in the lysates prior to immunoprecipitation.

### mSWI/SNF inhibition alters global protein synthesis

In order to investigate mSWI/SNF involvement in protein synthesis, we took advantage of available small molecule bromodomain inhibitors targeting BRD9 (iBRD9), BRD9 and BRD7 (TP472), and SMARCA2, SMARCA4 and PB1 (PFI-3) (Fig. 3A). We treated HEK293T cells with these inhibitors for 24 hours and assessed protein synthesis for the last 30 minutes with puromycin or hour with L-azidohomoalanine (AHA). These assays revealed a substantial decrease of total protein synthesis upon each drug treatment (Fig. 3B-C).

**Figure 3.**
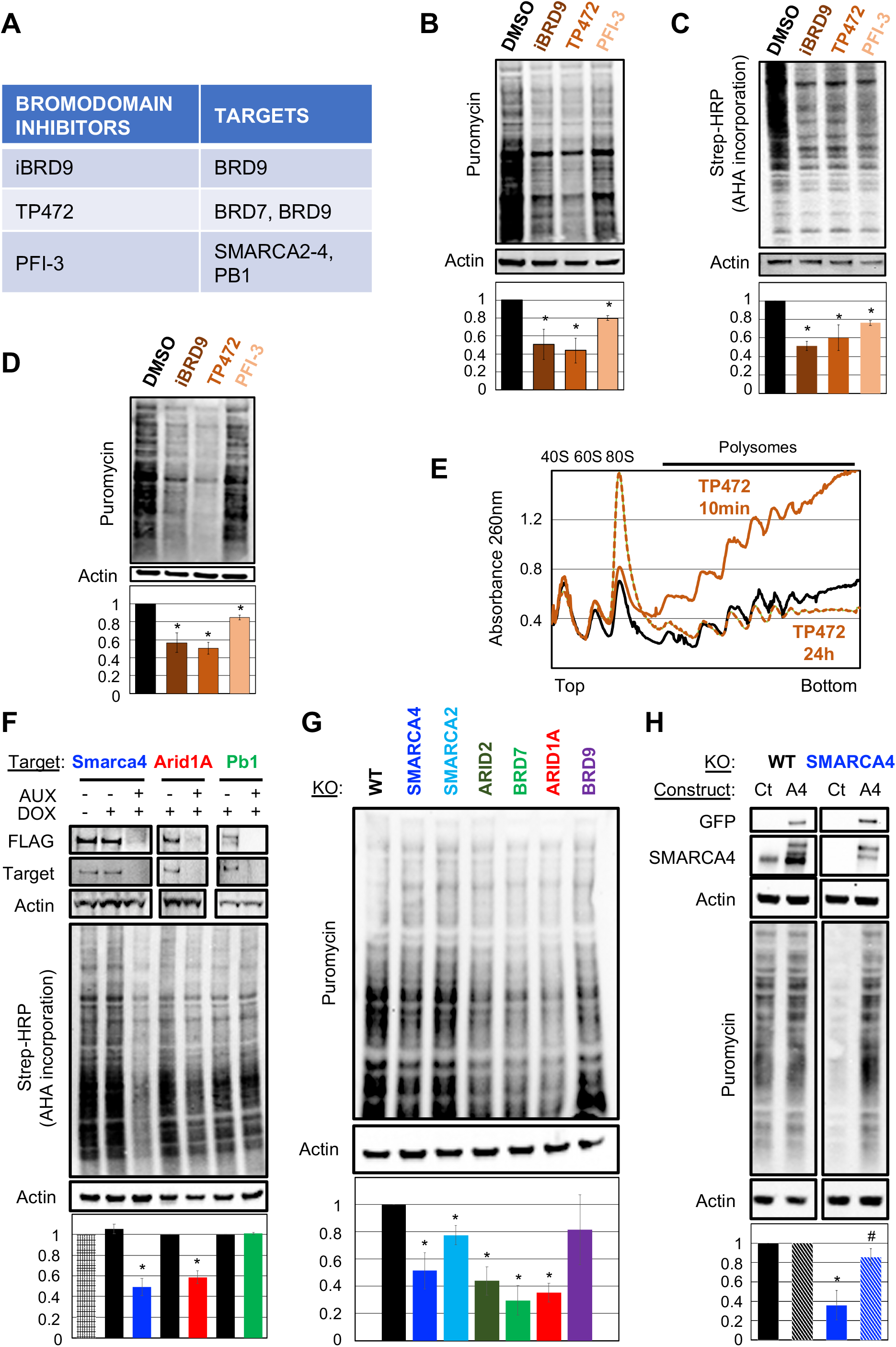
mSWI/SNF inhibition/depletion inhibits protein synthesis. **A.** mSWI/SNF bromodomain inhibitors and targets. **B.** Puromycin incorporation assays in HEK293T cells treated with 10μg/ml iBRD9, 5μM PFI-3, and 5μM TP472 for 24 hours. Puromycin was added for the last 30 minutes. Lower panel: quantification of three independent experiments. **C.** L-azidohomoalanine AHA incorporation assays in HEK293T cells treated as in A for 24 hours. AHA was added for the last hour. Lower panel: quantification of three independent experiments. **D.** Puromycin incorporation assays in HEK293T cells treated as in A for 10 minutes. Lower panel: quantification of three independent experiments. **E.** Representative polysome profiles from HEK293T cells treated with 5μM TP472 for 24 hours (dotted brown) and 10 minutes (brown). **F.** AHA incorporation in ES cells after 2 hours of auxin (AUX) inducible degradation of Smarca4, Arid1A and Pb1. Doxycycline (DOX) induces the expression of the Fbox degrading endogenous subunits tagged with FLAG and a degron interacting with the Fbox in the presence of auxin. Lower panel: quantification of three independent experiments. **G.** Puromycin incorporation assays in HAP1-WT and knock-out cells. Lower panel: quantification of three independent experiments. **H.** Puromycin incorporation assays in HAP1-WT and HAP knock-out for SMARCA4 after transient transfection of GFP-SMARCA4 or a control vector. Lower panel: quantification of three independent experiments. * two-tailed students’ t-test, p < 0.05.

We then wanted to decipher if the translational defects observed upon mSWI/SNF inhibition could be a consequence of direct roles in translation, as opposed to their nuclear roles on transcription. We shortened mSWI/SNF inhibitor treatments to 10 minutes before assessing *de novo* protein synthesis activity to capture any changes in translation before transcriptional changes would be observed at the protein level, as RNA transcription, export and translation take at least 20 minutes for small proteins (*17, 18*). Puromycin incorporation is reduced upon 10 minutes of inhibition to a similar extent as at 24 hours (Fig. 3D). To confirm these results at a single cell level, we performed Simultaneous Overview of tri-Molecule Biosynthesis (SOM_3_B) (*19*). This method allows for the simultaneous measurement of puromycin and 5-BromoUridine incorporation (BrU, assessing transcription) in addition to viability and other cellular features at a single-cell level. SOM_3_B further confirmed the observed decrease in protein synthesis in live cells upon 10 minutes of mSWI/SNF inhibition (Supplementary Fig. S4A-B). While we did not detect any changes in transcription (data not shown), 10 minutes is a short timeframe for BrU incorporation and as a consequence the detected signal was not reliable. Protein synthesis remains active during all interphase stages of the cell cycle and is reduced during mitosis (*20*). To identify if the observed inhibition is consistent as cells transit though the cell cycle, we assessed IdU incorporation (S phase) and expression of Cyclin B1 and phosphorylation of histone H3 on Serine 10 to assign each cell to its cell cycle phase (*19*). mSWI/SNF inhibition reduced puromycin incorporation across all stages of interphase and altered translation very modestly in mitotic cells (Supplementary Fig. S4C-D).

We then wanted to assess translational defects by polysome profiling assays. While after 24 hours of treatment each drug increased the levels of the 80S monosome and decreased the amount of polysomes, typical of a translation initiation defect, 10 minutes of mSWI/SNF inhibition led to increased polysomes, which could be reflective of ribosome collision (Fig. 3E and Supplementary Fig. S4E-H). Together, these data demonstrate that mSWI/SNF inhibition decreases global protein synthesis and that these defects are consistent with a direct function for mSWI/SNF subunits in translation.

### Depletion of specific mSWI/SNF subunits alters translation

To rule out any potential off-target effect of mSWI/SNF bromodomain inhibitors, we confirmed our findings using engineered mouse embryonic stem cells containing an auxin-dependent degron within *Smarca4*, *Arid1A* or *Pb1* alleles, allowing a complete degradation of endogenous protein upon auxin treatment (*21*). Addition of auxin for 2 hours inhibits *de novo* protein synthesis for Smarca4 and Arid1A but not Pb1 (Fig. 2E), highlighting the involvement of specific subunits or complexes in protein synthesis. Depletion of Smarca4 decreases protein synthesis by 50% but equally up” and downregulates expression of specific RNAs (Fig. 3F and (*21*)), validating potential independent roles for mSWI/SNF in translation in non-transformed cells.

In order to assess more mSWI/SNF subunits, we detected steady-state translational levels in HAP1 cells depleted for specific PBAF (*BRD7*, *ARID2*), BAF (*ARID1A*) and ncBAF (*BRD9*) subunits, as well as for the two ATPases (*SMARCA2* and *SMARCA4*) (Supplementary Fig. S4I). mSWI/SNF subunits’ subcellular localization in the HAP1 wild-type (WT) cells are comparable to the localization observed in HEK293T cells (Supplementary Fig. S4J). Furthermore, treatment of HAP1-WT with iBRD9 and TP472 for 10 minutes decreased overall protein synthesis (Supplementary Fig. S4K). Puromycin integration assays revealed that *SMARCA4*, *ARID2*, *BRD7* or *ARID1A*-depleted cells displayed a 50-70% decrease in translation, while *SMARCA2* or *BRD9*-depleted cells were less affected (Fig. 3G). In order to assess the type of alterations our mSWI/SNF subunits-depleted cells could exhibit, we performed polysome profiling assays. While each HAP1 knockout cell line demonstrated altered polysome profiles compared to WT cells, these profiles were distinct. *SMARCA4*, *ARID2* or *BRD7*-depleted cells showed identical profiles with an increase in initiating and early polysomes and decrease in late polysomes (Supplementary Fig. S5A-C); *SMARCA2* or *ARID1A*-depleted cells exhibited a decrease across the entire profile (Supplementary Fig. S5D-E); and *BRD9*-depleted cells, while being more variable, demonstrated a consistent increase in polysomes (Supplementary Fig. S5F-G). These distinct polysome profile alterations upon depletion of mSWI/SNF subunits from different complexes are consistent with a specificity of each complex in translation. As described in the nucleus, these results suggest different roles for BAF, PBAF and ncBAF in the cytoplasm. Finally, to demonstrate that altered translation is specific to the lack of SMARCA4 in *SMARCA4*-depleted cells we transiently transfected GFP-SMARCA4 in these cells. While the overexpression of SMARCA4 in HAP1-WT cells did not change levels of translation, expression in *SMARCA4*-depleted cells rescued the defects observed upon knock-out (Fig. 3H). Together, these results reveal that depletion of specific mSWI/SNF subunits decreases overall translation and that such roles are conserved in mice.

### mSWI/SNF inhibition and depletion sensitize cells to mTOR-PI3K inhibitors

Because mSWI/SNF inhibition decreases protein synthesis (Fig. 3B-D), we hypothesized that mSWI/SNF inhibitors could increase cells’ sensitivity to translation inhibitors. To assess this possibility, we performed cell viability assays using AZD8055 and BEZ235, inhibiting the kinases mTOR and mTOR-PI3K, respectively, in combination with mSWI/SNF bromodomain inhibitors. While 72 hours of iBRD9, TP472 or PFI-3 treatment alone decrease HEK293T viability, iBRD9 and TP472 decreased further the viability of cells co-treated with AZD8055 or BEZ235 (Fig. 4A-B).

**Figure 4.**
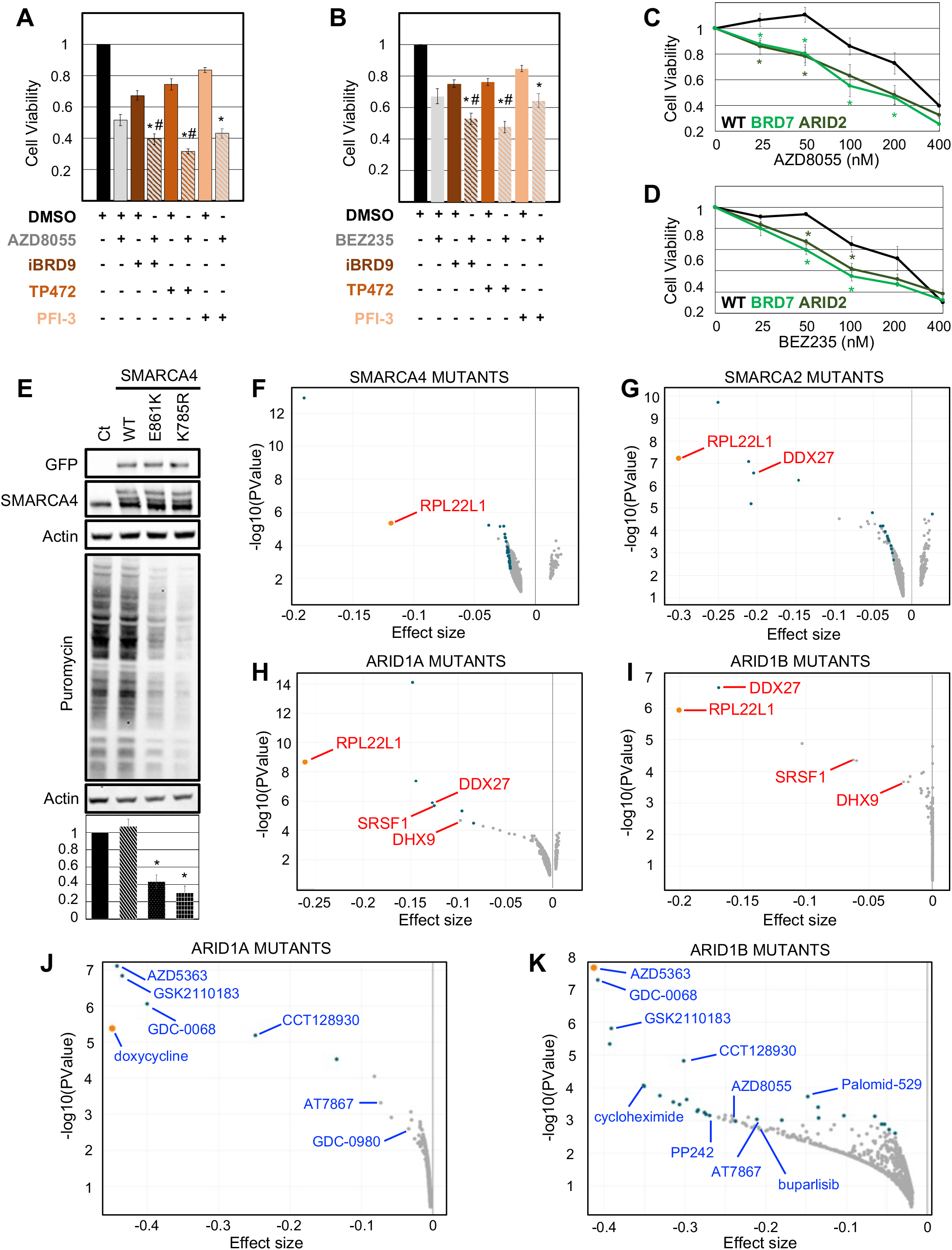
mSWI/SNF inhibition, depletion or pathogenic mutation sensitizes cells to translation pathway inhibitors. **A-B.** Cell viability assays in HEK293T cells treated with 10μg/ml BRD9, 5μM PFI-3, 5μM TP472, and 50nM AZD8055 (mTOR inhibitor, **A**) or 50nM BEZ235 (dual mTOR-PI3K inhibitor, **B**) for 72h. Averages of eight independent experiments are represented. * compared to its respective mSWI/SNF inhibitor alone, # compared to AZD8055 or BEZ235. **C-D.** Cell viability assays in HAP1-WT and PBAF subunits knockout cells treated with 25-400nM AZD8055 (**C**) or 25-400nM BEZ235 (**B**) for 72h. Averages of six (AZD8055) and five (BEZ235) independent experiments are represented. * compared to WT. **E.** Puromycin incorporation assays in HAP1-WT cells transiently overexpressing GFP-SMARCA4 ATPase domain mutations. Lower panel: quantification of three independent experiments. **F-I.** Genetic dependency of SMARCA4 (**F**), SMARCA2 (**G**), ARID1A (**H**) and ARID1B (**I**) mutated cancer cells compared to all other cancer cells in the DepMap database (combined RNAi dataset). **J-K.** Drug sensitivity of ARID1A (**J**) and ARID1B (**K**) mutated cancer cells compared to all other cancer cells in the DepMap database (PRISM repurposing primary screen 19Q4 dataset). Orange dot: most prominent translation-related gene/drug. Blue dots: q value <= 0.05. * and # two-tailed students’ t-test, p < 0.05.

Based on these results we further hypothesized that the depletion of mSWI/SNF subunits would sensitize cells to translation inhibition. Cell viability assays in knockout HAP1 cells revealed that the depletion of *ARID2* and *BRD7* increased sensitivity to a range of concentrations of AZD8055 and BEZ235 (Fig. 4C-D), while depletion of *SMARCA2*, *SMARCA4*, *ARID1A* or *BRD9* did not (Supplementary Fig. S6A-D). Together, these data demonstrate that mSWI/SNF inhibition and depletion of specific subunits creates a vulnerability to mTOR-PI3K inhibition in a variety of cell types.

### SMARCA4 pathogenic mutations decrease protein synthesis

mSWI/SNF subunits are frequently mutated in cancers and most of these mutations are considered loss-of-function (*1–3*). Furthermore, some heterozygous missense mutations act in a dominant-negative fashion (*22*). Therefore, we hypothesized that missense mutations of mSWI/SNF subunits could alter translation. To test this hypothesis, we took advantage of two well characterized *SMARCA4* pathogenic mutations: p.K785R, located in the ATP cleft and acting as a catalytic mutant, and p.E861K, located in the DNA groove and acting to disrupt DNA binding (*22*). We transiently overexpressed these mutations in HAP1-WT cells and assessed protein synthesis by puromycin integration assays. Overexpression of either mutation decreased overall translation (Fig. 4E). Therefore, these alterations act as dominant-negative mutants and SMARCA4’s catalytic activity and DNA groove are both required for normal translation.

### Cancer cells harboring mSWI/SNF mutations exhibit a genetic dependency on translation factors

We further hypothesized that mSWI/SNF mutated cells could be dependent on translation. In order to test this hypothesis, we took advantage of the genetic dependency study of the DepMap database reporting RNAi against 17,309 genes in 712 cell lines (*23*). Fourteen mSWI/SNF subunit pathogenic mutations were dependent on one or more translation factors when compared to all other cell lines in the dataset (Fig. 4F-I and Supplementary Fig. S6E-N). In accordance with decreased translation associated with *SMARCA4* mutations (Fig. 4E), *SMARCA4*-mutated cancer cells demonstrated a genetic dependency on the ribosomal subunit RPL22L1 and the RNA helicase DDX27 (Fig. 4F). *SMARCA2*-mutated cancer cells also exhibited a dependency on RPL22L1 (Fig. 4G), as well as *ARID1A* and *ARID1B*-mutated cancer cells (Fig. 4H-I). Cancer cells harboring mutations in either of these two BAF subunits also showed dependency on the RNA helicases DDX27, DDX9 and SRSF1, a splicing factor with a direct role in translation (Fig. 4H-I).

### mSWI/SNF mutations sensitize cancer cells to translation pathway inhibitors

The specific genetic dependency of *SMARCA2*, *SMARCA4*, *ARID1A* and *ARID1B*-mutant cancer cell lines led us to speculate that these mutations could sensitize cancer cells to translation inhibitors. We analyzed the PRISM repurposing primary drug sensitivity screen in the DepMap database testing 578 cell lines with 4,686 compounds (*24*). Cancer cells mutated for either ATPase did not present any strong sensitivity to specific inhibitors when compared to all other cells in the database (q value < 0.05) but did sensitize cells to some translation pathways inhibitors (Supplementary Fig. S7A-B). However, cancer cells harboring *ARID1A* or *ARID1B* mutations were primarily more sensitive to inhibitors targeting the translation pathway: AKT, mTOR and PI3K inhibitors, as well as drugs directly affecting translation such as doxycycline and cycloheximide (Fig. 4J-K).

## Discussion

In the present study we demonstrate the substantial cytoplasmic presence of the mSWI/SNF subunits and their implications in protein synthesis. Decreased translation upon mSWI/SNF bromodomain inhibitor treatment suggests that these domains are necessary for mSWI/SNF targeting to the translation machinery. This is a strong possibility as bromodomains recognize acetylation, and the ribosome is highly acetylated. While both short and long-term mSWI/SNF inhibition leads to decreased translation to the same extent, polysome profile alterations are vastly different. These differences could reflect the mechanisms behind such inhibition: ribosome collision stalls elongation, leading to increased polysome density before triggering a global translational dysregulation by inhibiting initiation (*25*). Future studies will need to decipher if and how defects in mSWI/SNF subunits, which are extensively present within the initiating fractions, lead to global ribosome collisions.

While the knockout of each tested subunit decreased translation, cell sensitivity to mTOR-PI3K inhibition was increased only upon depletion of PBAF subunits. This could reflect specific roles in translation for different mSWI/SNF complexes. Indeed, a global role in translation does not preclude an additional role on specific mRNAs. For example, upon stress, the translation initiation factor eIF2α reduces global protein synthesis and preferentially translates specific mRNAs (*26*). Confirming potential specific roles, PBAF-depleted cell lines presented identical alterations in polysome profiles. Of note, polysome profiles from *SMARCA4* knockout resembled PBAF depletion and *SMARCA2* knockout resembled BAF (*ARID1A*); it will be interesting to further determine if cytoplasmic complexes would involve specific ATPases, in contrast to the nucleus where SMARCA2 and SMARCA4 are not complex-specific (*4*).

Analyses of the DepMap database revealed the genetic dependency of mSWI/SNF-mutated cancers and genetic dependency to translation factors and their specific sensitivity to translation pathway inhibitors. This could be used as a clinical strategy, such as PARP inhibitors for BRCA-deficient cancers (*27*). However, some subunits such as *BRD9* did not reveal such dependency even though iBRD9 did decrease translation. For such subunits, only specific mutations such as in the bromodomain could lead to increased sensitivity to translation pathway inhibitors. Therefore, such specificity would be missed if considering all mutations; bromodomain mutations are only a subset of *BRD9* pathogenic mutations. A systematic study of the consequences of each subunit’s cancer alterations will be necessary to assess consequences on translation and sensitivity to translation pathway inhibitors.

Large scale sequencing studies revealed the co-occurrence of activated mutations in PI3K and inactivating mutations of the mSWI/SNF complexes (*1, 3*). A cooperative mechanism between these two altered pathways has therefore been suggested (*1, 3*). While biologically this co-occurrence is not understood, the roles of mSWI/SNF in translation could be relevant. PI3K activation promotes protein synthesis and cellular proliferation (*28*). The decreased translation due to inactivating mutations in mSWI/SNF subunits could need the counteraction of increased translation through PI3K activating or *PTEN* inactivating mutations (*3*). Altering the translation of specific mRNAs is another route to enable a selective advantage to these cancer cells; moreover, these possibilities are not mutually exclusive. We do not observe any specific association between PI3K mutation and mSWI/SNF alteration in expression and localization in our breast cancer cases (data not shown). SMARCA4 was overexpressed in more than 90% of our cases, confirming the fact that SMARCA4 is likely an oncogene and that its low expression could be a favorable outcome in breast cancer (*3, 15*). On the contrary, SMARCA2 expression was extensively lost. It would be interesting to determine if such breast cancer cases would be more sensitive to translation inhibitors, especially when there is no SMARCA4 overexpression and/or when *SMARCA4* is mutated. Further, a proportion of breast cancer cases exhibited loss of PBAF subunits expression which could sensitize to translation pathways inhibitors. However, we also observed several cases with exclusive gain in cytoplasmic expression or opposite changes within different compartments. The consequences of such differences in subcellular functions could have great potential for the discovery of therapeutically targetable events.

## Methods

### Cell culture and drug treatments

HEK293T cells were obtained from ATCC. HAP1 and HAP1 SMARCA2 (HZGHC004055c012), SMARCA4 (HZGHC002878c004), ARID2 (HZGHC000907c009), ARID1A (HZGHC000618c010), BRD7 (HZGHC000923c010), BRD9 (HZGHC000934c010) knockout cell lines were obtained from and characterized by Horizon Discovery. Culturing media and media supplements were used based on supplier’s instructions. HAP1 cell lines were used under 30 passages. Mouse embryonic stem cells genome editing and culture are described in (*21*). BEZ235 (Apexbio), AZD8055 (AChemBlock) iBRD9 (Tocris), TP472 (Tocris) and PFI-3 (Tocris) inhibitors were used at the indicated concentrations on cells seeded since 24 to 72 hours.

### Antibodies

LDH (Santa Cruz, sc-133123), Streptavidin-HRP (Invitrogen SA100-01), Puromycin (Sigma, MABE343), Flag-tag (Sigma, F3165-.2MG), Veriblot IgG (Abcam, ab131366), Normal rabbit IgG (Cell signaling, 2729S), eIF3A (Abcam, ab118357), eIF4A1 (Abcam, ab31217), PABP (Abcam, ab21060), RPS6 (Abcam, ab58350), RPS3 (Abcam, ab140676), Beta actin (Abcam, ab8226), SMARCC2 (Abcam, ab71907), SMARCA4 (Abcam, ab110641; IP,WB), SMARCA4 (Abcam, ab108318; IHC), SMARCC1 (Abcam, ab172638), SMARCA2 (Abcam, ab15597; WB), SMARCA2 (Sigma HPA029981; IHC), PHF10 (Abcam, ab154637), PB1 (Abcam, ab196022; WB, IHC), BRD7 (Abcam, ab56036; WB), BRD7 (Protein tech 51009-9-AP; IHC), DPF2 (Abcam, ab134942), ARID2 (Abcam, ab51019; WB), ARID2 (Abcam ab113283; IHC), ARID1B (Abcam, ab84461), ARID1A (Cell signaling, 12354S), BRD9 (Bethyl laboratories, A303-781A), GLTSCR1 (Santa Cruz, sc-515086), SS18 (Cell signaling, 21792S), SMARCB1 (Abcam, ab192864), Cyclin B1 (Fisher Scientific, BDB554177), Phospho-Ser28-Histone3 (BioLegend, 641002).

### Cell Fractionation

Cell fractionation was performed as in (*29*). For the western-blot analysis a comparable fraction of each compartment was loaded on a gel.

### Immunohistochemistry

SMARCA4 was performed on a Roche Ventana Benchmark Ultra instrument (Tucson, AZ) using CC1 retrieval and antibody dilution at 1:25. SMARCA2 (1:300), BRD7(1:400) and PB1 (:2000) were performed manually using Tris-EDTA antigen retrieval (pH 9). ARID2 (1:250) was performed manually using citrate antigen retrieval (pH 6).

### Immunoprecipitation

Immunoprecipitation was performed as in (*16*).

### Translation assays

L-azidohomoalanine (AHA) incorporation were performed as in (*16*). Puromycin incorporation assays were performed as in (*30*). Antibodies intensities from western blot membranes were analyzed using ImageJ. Because of the string smear obtained for these western blots, loading controls (actin) were ran on separate membranes.

### Polysome profiling

Polysome profiling were performed as in (*16*). Because of the high number of proteins tested and size restriction, identical fractions from polysome profiles were assessed with several gels.

### Viability assays

Viability assays were performed following supplier’s instructions from the CellTiter-Glo 2.0 Cell Viability Assay (Promega). Luminescence reading was performed using GloMax® Discover Microplate Reader (Promega).

### Simultaneous Overview of tri-Molecule Biosynthesis (SOM_3_B)

SOM_3_B was performed as described previously in (*19*). Live cells were treated with inhibitors for 10 minutes and spiked with IdU, BrU. Single cell mass-cytometry datasets generated and presented are available in flowrepository.org with the following identifier: *public identifier will be generated at time of publication*.

### Quantification and Statistical analysis

Please refer to the figure legends or the experimental details for description of sample size (n) and statistical details. Cell culture assays were performed at least three independent times. Data are expressed as mean ± SEM (Standard Error of the Mean). Differences were analyzed by two-tailed Student’s t test, one-way ANOVA, P-values ≤ 0.05 were considered significant and annotated with *.

## Acknowledgments

The authors would like to thank Drs. Joshua Back and Johnathan Whetstine for their helpful discussions of the data, and Ellen Gomulia and Ivy Mangonon for assistance with SMARCA4 immunohistochemistry. G.R.C. is supported by the Howard Hughes Medical Institute, grants from the NIH, CA163915 and NS046789 and a grant from the Department of Defense, Breast Cancer Research fund. C.M.W. was supported by a grant from GSK, Sir James Black Fellowship. S.C.K and S.C.B received support from the NIH: T32GM007276 (S.C.K), 1DP2OD022550-01 (S.C.B.), 1R01AG056287-01 (S.C.B.), 1R01AG057915-01 (S.C.B.) and 1U24CA224309-01 (S.C.B.).

