## Supplemental Figures for "mSWI/SNF interacts with the ribosome and its inhibition/mutations alter translation and sensitize to mTOR/PI3K inhibitors"

### Supplementary Figures and Legends

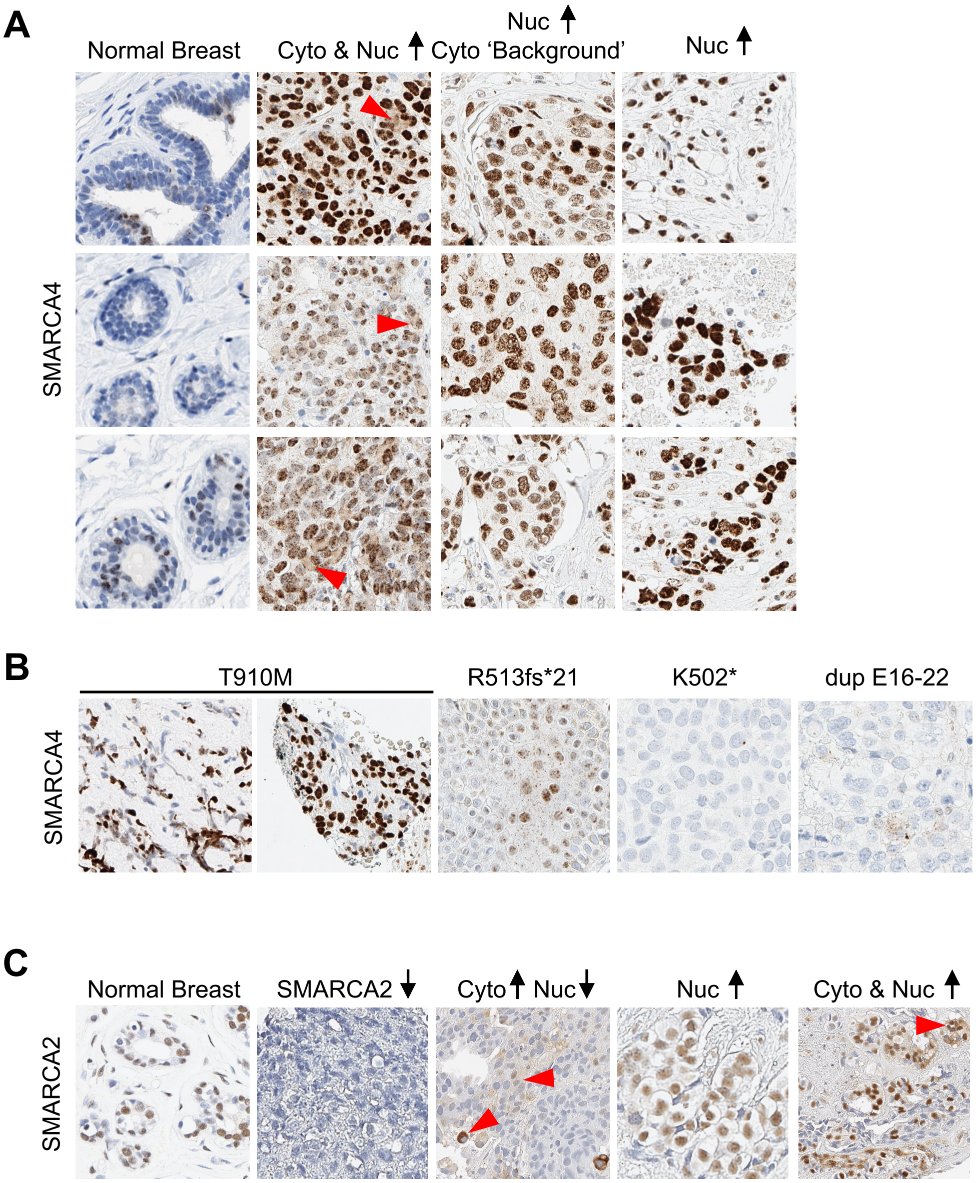


***Figure S1 – SMARCA2 and SMARCA4 expression and subcellular localization are altered in breast cancers.*** *Representative examples of immunohistochemistry from breast cancer cases. SMARCA4 expression in nine wild-type tumors (**A**) and tumors with known SMARCA4 pathogenic mutations (**B**). SMARCA2 expression in four breast cancer cases (**C**). 20X images. Cyto = cytoplasmic, Nuc = nuclear.*

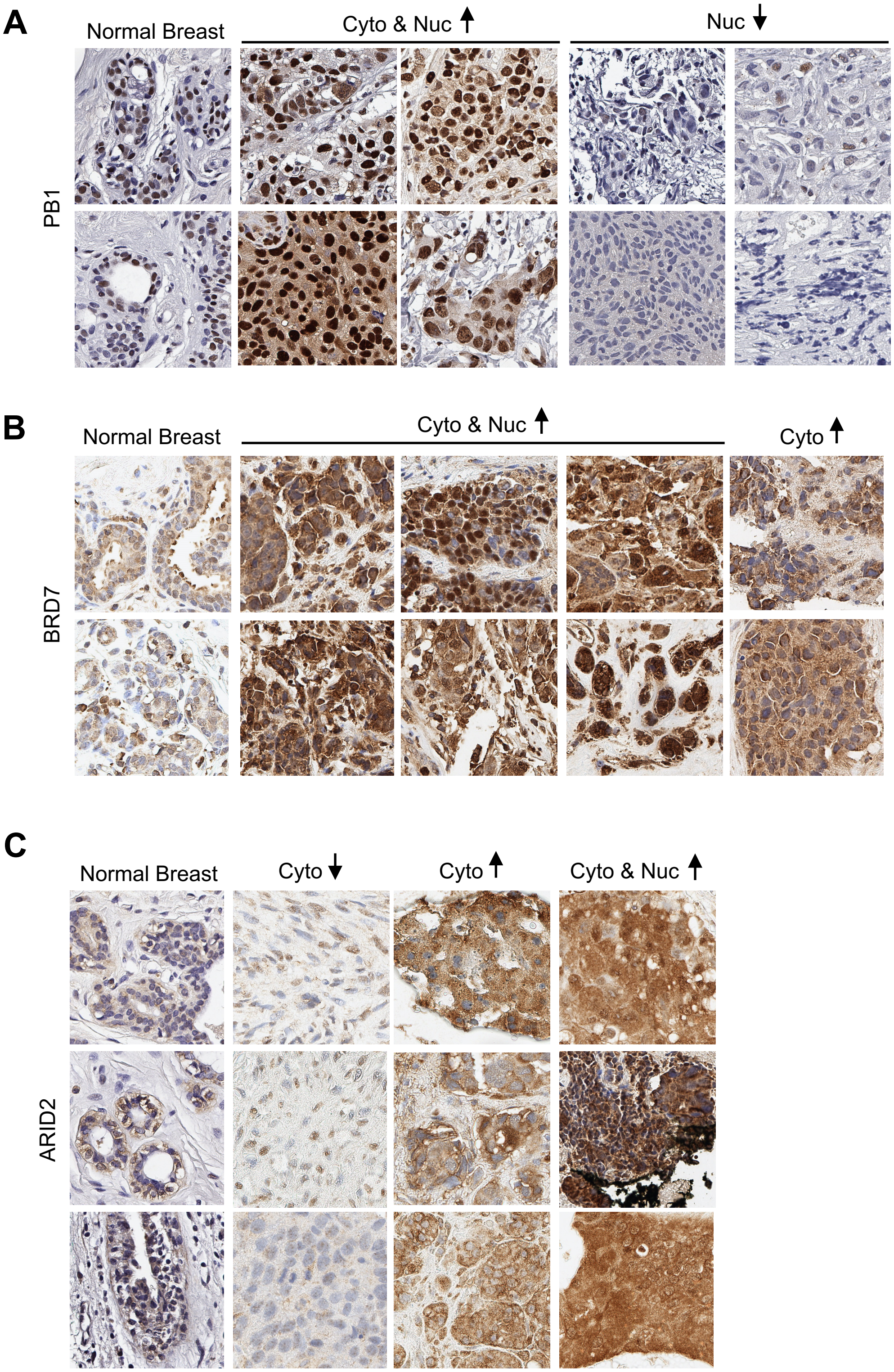


***Figure S2 – PB1, BRD7 and ARID2 expression and subcellular localization are altered in breast cancers.*** *Representative examples of immunohistochemistry from breast cancer specimens for PB1 (**A**, eight cases), BDR7 (**B**, eight cases) and ARID2 (**C**, nine cases). 20X images. Cyto = cytoplasmic, Nuc = nuclear.*

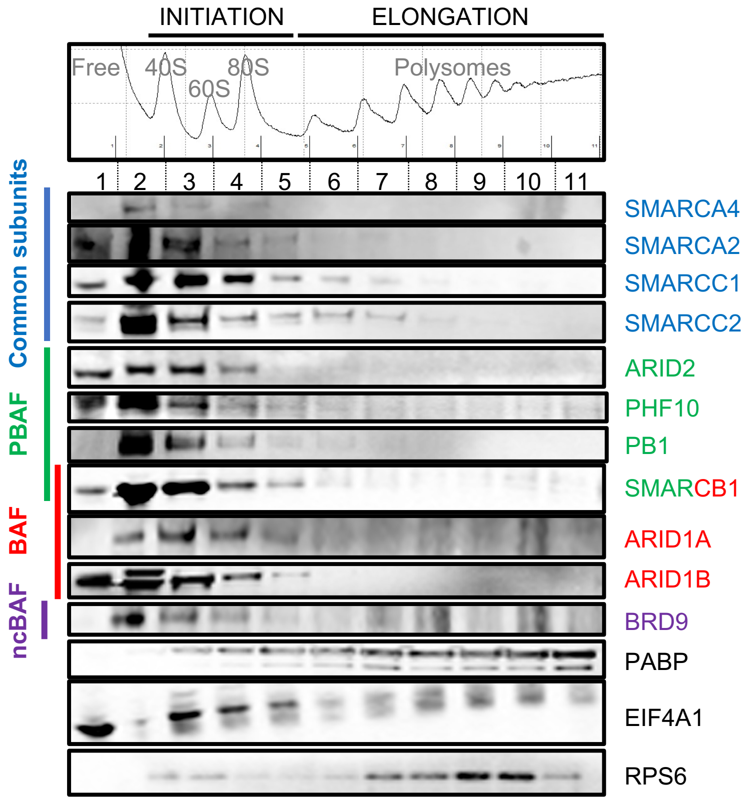


***Figure S3 – mSWI/SNF subunits localize in the cytoplasm and interact with the translation machinery.*** *Immunoblots from HEK293T polysome profiles fractions.*

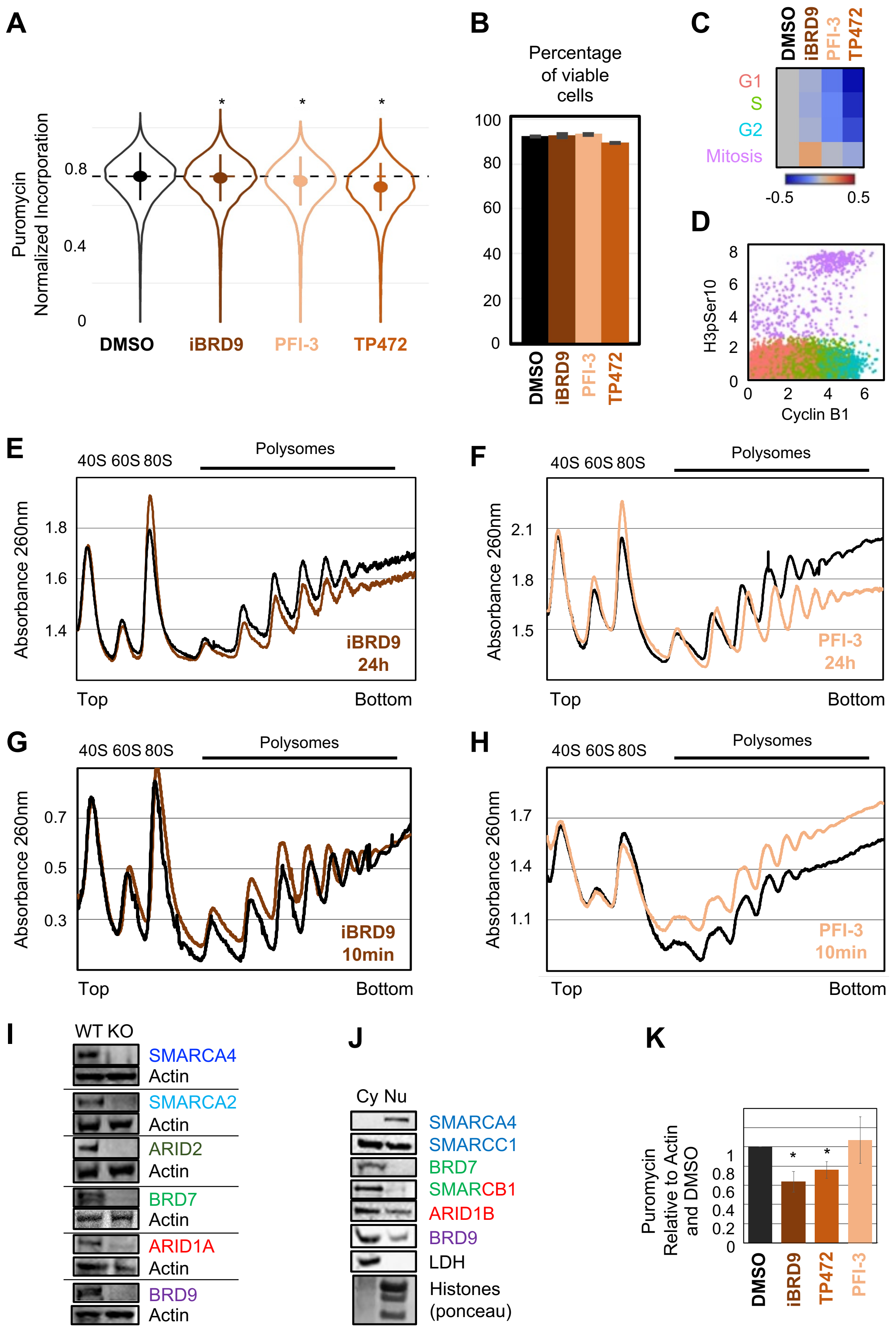


***Figure S4.*** *Legend on next page.*

***Figure S4 – mSWI/SNF inhibition/depletion inhibits protein synthesis. B.*** *SOM$_3$B analysis of HEK293Tcells treated with 10μg/ml iBRD9, 5μM PFI-3 or 5μM TP472 for 10 minutes, from three independent experiments. * ANOVA, $p < 2.2e-16$.* ***B.*** *Percentage of viable cells from SOM$_3$B analysis reported in Fig. 2C. The average of three independent experiments are represented.* ***C.*** *Puromycin integration across different phases of the cell cycle in single cells detected by SOM$_3$B analysis. The results from three independent experiments are represented as Log2 puromycin integration compared to DMSO.* ***D.*** *Detection of H3pSer10 and Cyclin B1 by SOM$_3$B analysis. The colors correspond to colors of the cell cycle phases represented in D.* ***E****. Representative polysome profiles from HEK293T cells treated with 10μg/ml iBRD9 for 24 hours (brown).* ***F****. Representative polysome profiles from HEK293T cells treated with 5μM PFI-3 for 24 hours (beige).* ***G****. Representative polysome profiles from HEK293T cells treated with 10μg/ml iBRD9 for 10 minutes (brown).* ***H****. Representative polysome profiles from HEK293T cells treated with 5μM PFI-3 for 10 minutes (beige).* ***I.*** *Immunoblot from HAP1-WT and knockout cells.* ***J.*** *Immunoblot from fractionated HAP1-WT cells (Cy: cytoplasm, Nu: nucleus and chromatin combined).* ***K.*** *Puromycin incorporation assays in HAP1-WT treated with 10μg/ml iBRD9, 5μM PFI-3 or 5μM TP472 for 10 minutes. The quantification of four independent experiments is represented. * two-tailed students' t-test, $p < 0.05$.*

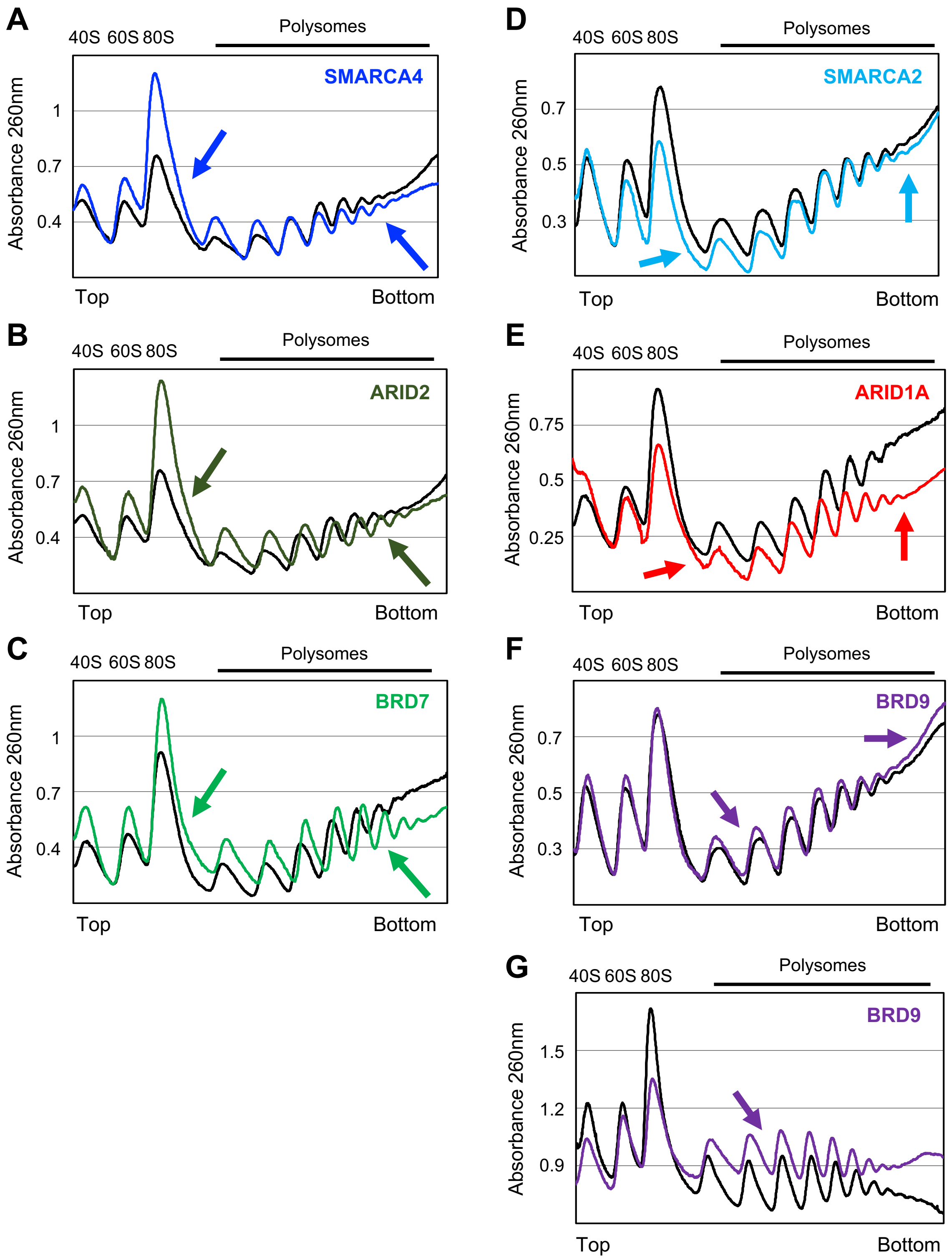


***Figure S5 – mSWI/SNF knockout cells exhibit specific polysome profile defects.*** *Representative traces of polysome profiles from HAP1 knockout cells (black lines: WT).*

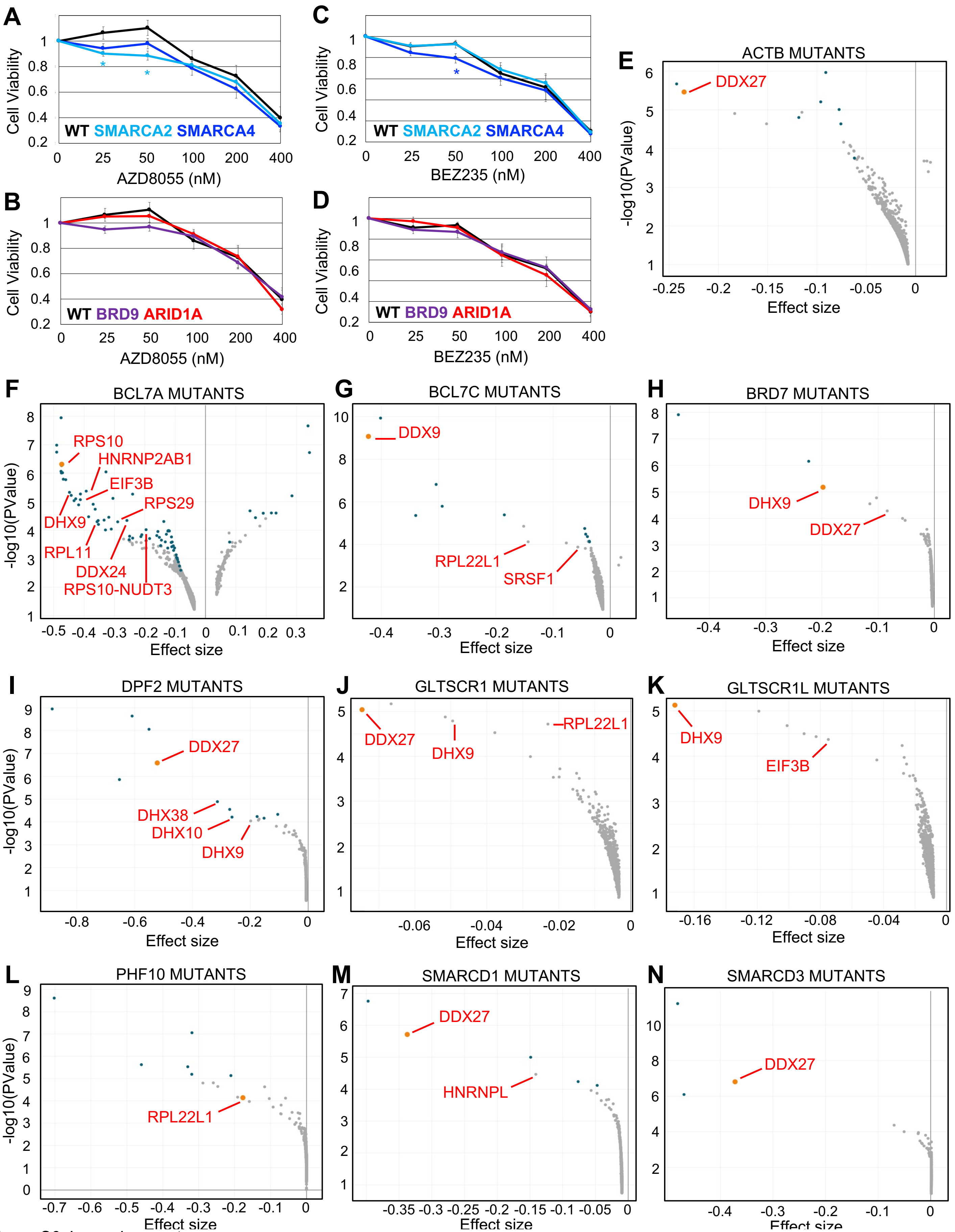


***Figure S6.*** *Legend on next page.*

***Figure S6 – mSWI/SNF inhibition, depletion or pathogenic mutation sensitizes cells to translation pathway inhibitors. A-D.*** *Cell viability assays in HAP1-WT and ATPases (blue), BAF (red) or ncBAF (purple) subunit knockout cells treated with 25-400nM AZD8055 (**A-B**) or 25-400nM BEZ235 (**C-D**) for 72h. Averages of six (AZD8055) and five (BEZ235) independent experiments are represented. * compared to WT. # two-tailed students' t-test, p < 0.05.* ***E-N.*** *Genetic dependency of ACTB (**E**), BCL7A (**F**), BCL7C (**G**), BRD7 (**H**), DPF2 (**I**), GLTSCR1 (**J**), GLTSCR1L (**K**), PHF10 (**L**), SMARCD1 (**M**) and SMARCD3 (**N**) mutated cancer cells compared to all other cancer cells in the DepMap database (combined RNAi dataset). Orange dot: most prominent translation-related gene. Blue dots: q value <= 0.05.*

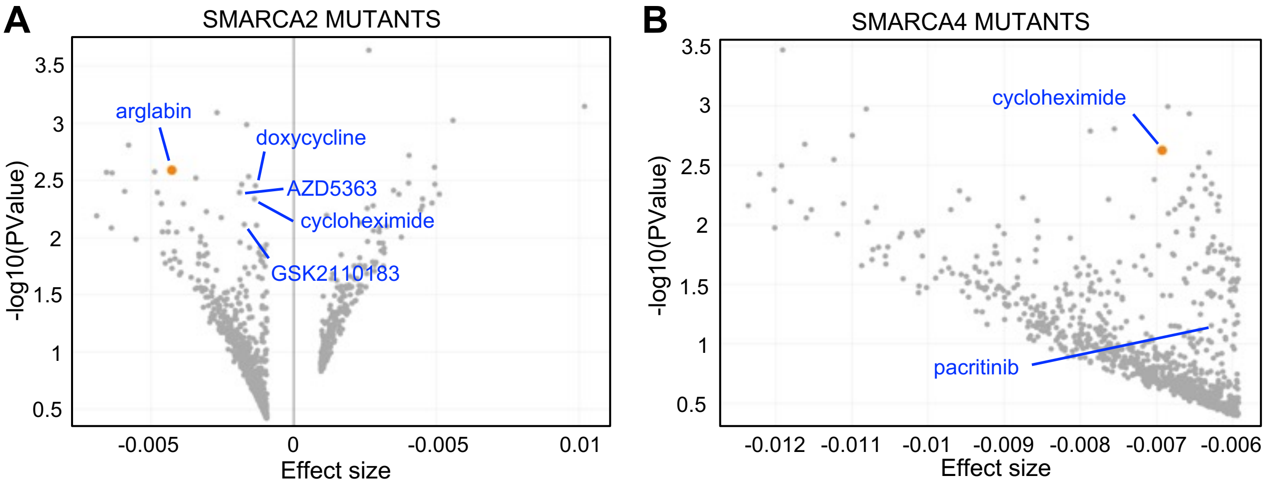


***Figure S7 – mSWI/SNF pathogenic mutations sensitize cells to translation pathway inhibitors.*** *Drug sensitivity of SMARCA2 (**A**) and SMARCA4 (**B**) mutated cancer cells compared to all other cancer cells in the DepMap database (PRISM repurposing primary screen 19Q4 dataset). Orange dot: most prominent translation-related drug. Blue dots: q value <= 0.05.*
